# Toward Reliable Synthetic Omics: Statistical Distances for Generative Models Evaluation

**DOI:** 10.1101/2025.05.08.652855

**Authors:** Raffaele Marchesi, Nicolò Lazzaro, Gianluca Leonardi, Federica Rignanese, Stefano Bovo, Marco Chierici, Giuseppe Jurman

**Affiliations:** Department of Mathematics, University of Pavia, Via Ferrata 5, 27100, Pavia, Italy; Data Science for Health, Fondazione Bruno Kessler, Via Sommarive 18, 38123, Trento, Italy; Department of Cellular, Computational and Integrative Biology - CIBIO, University of Trento, Via Sommarive 9, 38123, Trento, Italy; Department Biomedical Sciences, Humanitas University, Via Rita Levi Montalcini 4, 20072, Pieve Emanuele (MI), Italy

**Author notes:** These authors contributed equally to this work as co-first authors (Raffaele Marchesi, Nicolò Lazzaro).

**Keywords:** Statistical distances, Artificial intelligence, Cancer, Transcriptomics

## Abstract

**Background:** Synthetic data generation is emerging as an approach to overcome the limitations of real-world data scarcity in *omics* studies, especially in precision medicine and oncology. *Omics* datasets, with their high dimensionality and relatively small sample sizes, often lead to overfitting, especially in deep learning models. Generative models offer a promising way to generate realistic synthetic data preserving the original data distribution. However, there is still no objective consensus on how to evaluate their performance. In this study, we set out to validate generative networks for transcriptomics data generation by using statistical distances as robust evaluation metrics.

**Results:** We observe that statistical distances enable simultaneous evaluation of global and local data fidelity of generated synthetic data. Because these distances satisfy the properties of true metrics, they also enable formal hypothesis testing to assess whether generative models have in fact converged or are merely approaching the reference distribution. Crucially, optimizing for these distances was found to implicitly select models maximizing other widely used metrics of generative performance, providing evidence of their broad applicability. Overall, our findings indicate that the adoption of these metrics can play a key role in guiding the development of generative models across a wide range of domains.

## Introduction

*Omics* data are becoming everyday more relevant in precision medicine. In particular, RNA sequencing (RNA-seq) has become a milestone technique in precision oncology (Wang et al., 2009; Lau et al., 2019), providing insights into the underlying mechanisms of cancer development, progression, and response to therapy. Transcriptomics data enabled the study of how gene expression changes in different organisms with high sensitivity and low bias while having scalable capabilities for high-throughput applications (Mortazavi et al., 2008; Su et al., 2014). Despite the increasingly central role of this type of data for research and clinical purposes, their usage presents several challenges. In fact, the information retrieved from this technology is characterized by both high dimensionality and nonlinear expression patterns, making them complex to manage and analyze. Furthermore, this type of data is characterized by limitations regarding its use, as it is regulated by restrictive privacy constrains and concerning accessibility since not many clinical and research facilities have the appropriate instrumentation. These aspects have an impact on the quality of research and innovation by lengthening the study timeline and slowing the development of new solutions or treatments. Under these circumstances, synthetic data can represent a valuable solution. This new approach aims to recreate data mimicking real-world patient data satisfying privacy regulations and preserving quality while improving accessibility by limiting the sparsity and allowing data augmentation. In cases where data can be skewed or underrepresented, synthetic data generation can be utilized as a mitigation methodology. Moreover, synthetic data can be used to improve the performance of AI models in terms of generalization and effectiveness (Frid-Adar et al., 2018; Wong et al., 2020). Although synthetic data generation is a growing branch of research, there is no consolidated and shared procedure for their calculation and evaluation. Several deep learning architectures could be used for synthetic data generation. Variational Autoencoders (VAEs) and Generative Adversarial Networks (GANs) are among the key methodologies currently used for generating synthetic tabular data in healthcare (Micheletti et al., 2023; Liu et al., 2025; Jadon and Kumar, 2023). In the same domain, Diffusion Models (DM) have been successfully used to generate high-resolution, realistic images (Han et al., 2023; Pozzi et al., 2024) and now are emerging as a tool to generate synthetic tabular data (Kotelnikov et al., 2022; Villaizán-Vallelado et al., 2024), including transcriptomics (Lacan et al., 2024). To assess the goodness of data generated with these models, several metrics have been proposed (Yale et al., 2020; Murtaza et al., 2023; Borji, 2022; Hernadez et al., 2022).

This work aims to validate generative networks for data generation and to propose two statistical distances as evaluation metrics: the energy distance and the pointwise empirical distance. These distances make it possible to directly evaluate the global distribution of real and synthetic data at once, whilst still allowing for clustering approaches and local evaluation.

## Methods

### Statistical Distances

Statistical distances offer a simple and robust approach to compare the similarity between probability distributions or sets of empirical data points. In the context of generative models, a distance serves as a quantitative measure of how well a synthetic dataset approximates the real data distribution. Distances that remain valid in high-dimensional spaces, such as those encountered in transcriptomics, are especially relevant in assessing the fidelity and diversity of generative models. Moreover, when these distances satisfy the properties of a true metric, they enable additional analyses such as hypothesis testing on equality of distributions.

**Energy Distance** (Rizzo and Székely, 2016) is a nonparametric measure that stands out for its balance between tractability and interpretability. Given two random variables *X* and *Y* taking values in a normed space *𝒳* with norm ∥ · ∥, the squared energy distance is defined as

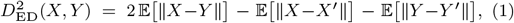

where *X*^′^ and *Y* ^′^ are i.i.d. copies of *X* and *Y*, respectively. The statistic *D*_ED_(*X, Y*) is zero if and only if *X* and *Y* follow the same distribution, making it a proper metric. A key strength of the energy distance is that it captures broad differences in distribution location and spread without requiring explicit modeling of the underlying data-generating process. It also remains sensitive to distributional mismatches in high dimensions, making it suitable for *omics* datasets. Furthermore, its low computational cost (O(*n*^2^) in the sample size) and differentiability make it a valid candidate for future direct integrations into neural networks penalty components to promote generalizability and avoid mode collapse.

**Pointwise Empirical Distance** (PED) (Lazzaro et al., 2024) refines global comparisons by emphasizing local geometric alignments between two distributions instead of overall match. This measure was originally developed to accommodate single-cell transcriptomics datasets, where the data is very noisy and overall manifold recapitulation can be more desirable than exact statistical fidelity. Unlike the energy distance, which essentially evaluates the expected pairwise distance between samples from different datasets, the PED compares the pairwise distance distributions themselves. Roughly speaking, it aligns points in the two empirical distributions to minimize overall distortion and then measures how far apart those matched points remain.

Similarly to the energy distance, it compares i.i.d. pairs (*X, X*^′^) and (*Y, Y* ^′^) from the random variables *X* and *Y* in a normed space. Rather than simply averaging the norms of pointwise differences, however, it compares the induced distance profiles of each sample to those of its counterpart under an optimal alignment. Formally,

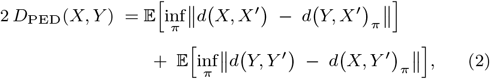

where *d*(·, ·) denotes the distance between two points in the chosen norm, and *π* ranges over all permutations that align the pairwise distances in a way that minimizes the overall discrepancy.

Consequently, while computationally more expensive than the energy distance (owing to the permutation search), the PED is robust to noise and can be leveraged to detect subtler geometric discrepancies in high-dimensional data, whilst still being computable in polynomial time. This heightened sensitivity makes it particularly useful for certain domains, such as *omics*, where the local arrangement of points often encodes critical structural information. The infimum over permutations also tends to filter out noise, while the energy distance averages it into its expectation. On the other hand, unlike the energy distance, the PED is not differentiable and may be less useful when the data is easily normalizable and relatively noise-free.

**Local and Balanced Distances** are a natural extension to both statistical distances, obtained by evaluating them on local neighborhoods within the dataset. This can be done either by partitioning the samples with a clustering algorithm or directly evaluating the local neighborhood induced by every individual sample and its k-nearest neighbors. This measure can be evaluated alongside the one derived from the global distance estimate by considering their product to get a balanced distance assessing both global alignment and local fidelity with equal importance. In high-dimensional *omics* data, such a strategy can help ensure that the generated samples not only match the overall distribution of the real data but also capture important local structures, such as tissue-specific patterns.

**Likelihood Test** can be used to assess whether there is sufficient evidence to reject the null hypothesis that the synthetic and real data share the same distribution. By bootstrapping a statistical distance on subsets of a validation set, one obtains a reference distribution of distances that would be expected among samples drawn from it. Comparing the observed distance between real and synthetic samples to this reference distribution yields a formal test of whether the model is generating samples within the true data distribution.

**Identity Test** is a necessary complement to the likelihood test and evaluates whether synthetic data is not only realistic, but also distinguishable from the training set. Specifically, if the null distribution of the training set and that of a synthetic dataset are indistinguishable, it may indicate that the model’s ability to generate realistic data has not generalized (or the model was over-trained, overfitting the examples).

### Evaluation Metrics

The complexity of high-dimensional spaces, such as tabular gene expression data, poses a significant challenge in the assessment of the quality of the generated samples. Unlike image generation and natural language models, where human inspection can provide valuable insights into the quality and coherence of the synthetic data, our domain lacks such intuitive interpretability. In the absence of a gold standard, the validation approach usually relies on an ensemble of metrics that together should capture diverse statistical properties to ensure alignment to the real data distribution.

In the context of synthetic data evaluation, **Precision** measures the proportion of generated samples that fall within the real data manifold, while conversely **Recall** indicates the proportion of real data within the synthetic manifold (Sajjadi et al., 2018; Kynkäänniemi et al., 2019). As the harmonic mean of the two, the **F1 score** combines these complementary aspects into a single metric, providing a balanced measure of generation quality. When data is generated conditionally to labels, the F1 score can be measured for each individual class. The mean F1 score per class indicates if the generative model preserves label-specific characteristics, or if it captures only the full distribution of the real data.

**Frechet inception distance** (FID) is an unsupervised metric initially developed in a computer vision domain (Heusel et al., 2017). It is used to evaluate how realistic and diverse a set of synthetic data is compared to a real one by computing the Wasserstein-2 distance between the generated and real data distributions in a lower dimensional feature space. For tabular data, these features can be extracted from the last activation layer of a pre-trained classifier.

**Adversarial accuracy** (AA) (Yale et al., 2020) assesses if generated data are sufficiently close to true data (thus making augmentation effective) and, at the same time, sufficiently different from true training data (thus avoiding potential privacy breaches). Informally, AA represents the performance of an adversarial 1-nearest neighbor classifier that discriminates between true and synthetic data. *AA* = 0 indicates that synthetic data perfectly resemble true data (useful augmentation at the cost of high privacy loss), while *AA* = 1 indicates that synthetic data can be easily distinguished from real data (no privacy loss but limited utility of augmentation). A good generative model should aim for *AA* ≈ 0.5.

**Classification accuracy** (ACC) is a utility measure in a downstream application of the synthetic data. To evaluate synthetic data quality in a supervised learning setting, a classifier is trained on synthetic data and evaluated on a separate real data test set. This metric provides insights into whether the synthetic data preserves the discriminative properties of the real data for further machine learning tasks.

### Generative Adversarial Networks

Generative adversarial networks (GANs) (Goodfellow et al., 2014) are a family of generative models consisting of two neural networks: a generator *G* and a discriminator *D*. The generator is trained to produce a realistic synthetic data sample from a noise vector *z* sampled from a latent distribution. The discriminator learns to distinguish real data *x* from data generated by *G*. The training is adversarial and is defined as a minimax optimization problem:

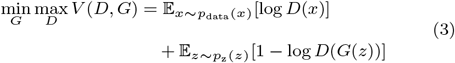

Adversarial training pushes the generator to produce more realistic samples, but also the discriminator to refine its classification. The result is the generation of progressively better synthetic data. However, this classical implementation of GANs is notoriously difficult to train because of mode collapse, vanishing gradients, and instability.

The Wasserstein GAN with gradient penalty (WGAN-GP) (Gulrajani et al., 2017) is designed to address these limitations. While the task of the discriminator of traditional GANs is a binary classification between real and fake samples, WGAN-GP introduces a Critic network that estimates the Wasserstein distance between real and synthetic data distributions. In addition, the gradient penalty term constrains the Critic’s gradient norm, enforcing a soft Lipschitz constraint on the network and penalizing large gradient magnitudes, thus preventing the Critic from becoming too sensitive to small changes in input. With this design, the WGAN-GP provides a more stable training, less prone to mode collapse.

The conditional version of WGAN-GP introduces conditioning labels *y* as input to both networks (Mirza and Osindero, 2014). The generator is trained to generate samples conditioned on specific attributes or metadata, and the discriminator can learn to distinguish samples based on these characteristics. This results in more control over the generation and gives a model that is able generate samples with desired characteristics. The losses of a conditional WGAN-GP are defined as:

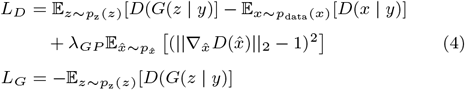

### Diffusion Models

Diffusion models are generative models that learn to reverse a multi-step noising process applied to data, enabling the generation of new samples from the learned data distribution (Sohl-Dickstein et al., 2015). In the forward process, Gaussian noise is incrementally added to data over multiple time steps, resulting in sequences of increasingly noisy data. In the reverse process, a neural network is trained to iteratively denoise these sequences, reconstructing data samples from noise. This approach has demonstrated state-of-the-art performance in various generative tasks, particularly image synthesis (Ho et al., 2020).

In the forward process, starting from an initial data sample *x*_0_, Gaussian noise is added over *T* time steps to produce a sequence *x*_1_, *x*_2_, …, *x*_*T*_. Each step can be defined as:

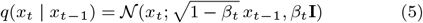

where *β*_*t*_ denotes the variance schedule controlling the noise level at each step, and *𝒩* represents a Gaussian distribution. This formulation ensures that as *t* approaches *T, x*_*T*_ approximates an isotropic Gaussian distribution (Sohl-Dickstein et al., 2015).

The reverse process aims to reconstruct *x*_0_ from *x*_*T*_ by iteratively removing the added noise. This is achieved by learning the conditional probability *p*_*θ*_(*x*_*t*−1_ | *x*_*t*_), parameterized by *θ*, which is modeled as:

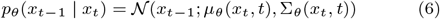

The covariance Σ_*θ*_ is usually fixed, and the model focuses on learning the mean *μ*_*θ*_:

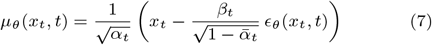

where *α*_*t*_ = 1 − *β*_*t*_ and 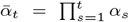. The term *ϵ*_*θ*_(*x*_*t*_, *t*) represents the model’s prediction of the added noise at step *t* (Ho et al., 2020). The model is trained to minimize the variational bound on the data likelihood, leading to the following objective:

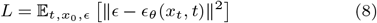

In this expression, *t* is uniformly sampled from {1, …, *T*}, *ϵ* ∼ *𝒩* (0, **I**), and *x*_*t*_ is obtained by adding noise to *x*_0_ according to the forward process. This loss function encourages the model to accurately predict the noise component at each step, facilitating effective denoising during generation (Ho et al., 2020).

Implementing diffusion models involves defining the forward and reverse processes, predicting the noise *ϵ*_*θ*_(*x*_*t*_, *t*) with a neural network, and minimizing the loss function. The neural network architecture often employed is a U-Net, which efficiently captures multi-scale features necessary for high-quality data synthesis. Training these models requires substantial computational resources due to the iterative nature of the denoising process and the need to model complex data distributions.

### Variational Autoencoders

The Variational Autoencoder (VAE) (Kingma and Welling, 2013) is a deep probabilistic generative model that learns a mapping from the probability distribution *p*_*θ*_(*x*) of the input data #x1D49F; = {*x*^(1)^, …, *x*^(*n*)^} to a lower-dimensional latent space that follows a simple probability distribution, like a standard Gaussian (i.e., *𝒩* (0, *I*)). This allows sampling a latent vector *z* from the latent distribution *p*_*θ*_(*z*) and generating new data points according to the conditional distribution *p*_*θ*_(*x*|*z*).

Generating a new sample would require knowing the true posterior distribution *p*_*θ*_(*z*|*x*), which cannot be computed due to the intractability of the marginal likelihood *p*_*θ*_(*x*). Variational inference is employed to approximate the true posterior *p*_*θ*_(*z*|*x*) with a simpler variational distribution *q*_*ϕ*_(*z*|*x*). This is achieved by employing an encoder network (also called an inference model) that maps the input data *x* to a distribution over the latent variables *z*, typically parameterized as a Gaussian with a mean *μ*(*x*) and variance *σ*(*x*). Similarly, a decoder (also called a generative model) reconstructs the original data from a latent variable *z*, sampled from the approximate posterior *q*_*ϕ*_(*z*|*x*), according to *p*_*θ*_(*x*|*z*).

The encoder and the decoder can be jointly optimized by maximizing a lower bound on the log-likelihood of the data, called the Evidence Lower Bound (ELBO), which is given by:

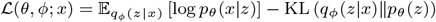

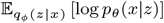 is the expected log-likelihood of the data given the latent variables, under the approximate posterior *q*_*ϕ*_(*z*|*x*). This is called reconstruction loss and encourages the decoder to generate data similar to *x* given the latent variables *z*. KL(*q*_*ϕ*_(*z*|*x*)∥*p*_*θ*_(*z*)) is the Kullback-Leibler (KL) divergence between the variational distribution *q*_*ϕ*_(*z*|*x*) and the prior *p*_*θ*_(*z*) and regularizes the latent space pushing it to be close to the simple prior distribution. To enable gradient-based optimization, the reparametrization trick is employed in order to make the sampling process differentiable.

A Conditional Variational Autoencoder (CVAE) (Sohn et al., 2015) is an extension of the standard VAE architecture in which both the encoder and the decoder are conditioned on some additional condition or label *y*. The model is trained by maximizing a modified version of the ELBO, which incorporates the conditional information provided by *y*:

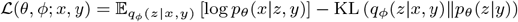

This allows the CVAE to generate samples that are not only faithful to the data distribution but also conditioned on the input label *y*.

### Datasets

The data used for the augmentation were extracted from two different public datasets. The first one is part of The Cancer Genome Atlas (TCGA), the most comprehensive public database on human genomics and transcriptomics that aims to increase the understanding of the genetic basis of a wide range of cancers. Specifically, the biospecimen repository of this dataset includes high-throughput genomic data from diseased and matched healthy samples spanning 33 cancer types (The Cancer Genome Atlas Research Network et al., 2013). We also considered the bulk gene expression dataset by the Genotype-Tissue Expression (GTEx) project (Lonsdale et al., 2013). This portal is a public resource created with data from three projects of the National Institutes of Health (NIH) to study the relationship between genetic variation and gene expression in normal human tissues and how changes contribute to common human diseases. The transcriptomic layer of TCGA (release date 2016-01-28) was retrieved using the RTCGA R package, while the eighth version (v8) with the TPM normalization was downloaded from GTEx. Data preprocessing, including duplicate removal, mapping of gene IDs, and standardization, was performed according to previous work (Lacan et al., 2024). Only genes included in the so-called L1000 landmark genes were used for both datasets. The L1000 genes were introduced by (Subramanian et al., 2017), which state that the expression of basically all remaining genes can be inferred by linear regression of those L1000, thus reducing the number of dimensions. Overall, the TCGA data in the present study consist of 10,156 patients, 978 landmark gene expression values, and 24 tissue types; the GTEx dataset includes 17,244 samples, 977 landmark gene expression values, and 26 tissue types. Data were split into training (TCGA: 8,124 samples; GTEx: 12,244 samples) and test (TCGA: 1,624; GTEx: 5,000 samples) sets. For some experiments, we also under-sampled the training set of both TCGA and GTEx datasets to 20% of the sample size (1,624 for TCGA and 2,448 for GTEx). This allowed us to evaluate the loss of quality in the synthetic data generated from a smaller training set, and it also gave us the remaining 80% of the samples to use as a validation set to measure the ability of the models to generalize outside the training distribution. The tissue types were used as covariates to condition the generative models.

### Experimental setup

We adopted a multi-stage process to evaluate the generative models on both the TCGA and GTEx datasets. First, each dataset was split into training and test sets in a stratified manner to preserve tissue-type proportions across the partitions. For TCGA, 8,124 samples formed the training set and 1,624 samples the test set; for GTEx, 12,244 samples were used for training and 5,000 for testing. Additionally, both datasets were under-sampled to create “reduced” versions validating the data efficiency and maximizing the statistical power of the likelihood and identity testing.

All generative models (GAN, VAE, and DM) were selected with a hyperparameter sweep strategy, exploring learning rates, network architectures, and other relevant parameters. The sweeps were performed with the Weights & Biases platform (Biewald, 2020) and each model was optimized either for a fixed number of epochs or until no appreciable improvement was observed. The selection of the best configuration for each model rested on minimizing the balanced statistical distances (a combination of global and local alignments, see Section “Statistical Distances”) between the synthetic and real data distributions on the training set.

Once trained, each model was used to generate synthetic datasets that mirrored the distribution of tissue types in the original data. We assessed the generated data on multiple quality metrics, including F1 Score, Adversarial Accuracy, and Frechet Inception Distance. In parallel, we evaluated the Balanced Energy Distance (ED) and Balanced Pointwise Empirical Distance (PED), both globally and locally, to verify whether the synthetic and real data aligned not only in their overall distribution but also in each tissue-specific subset. To further determine how faithfully the models captured the underlying data generation process, we conducted a likelihood test (measuring whether synthetic data could plausibly come from the real-data distribution) and an identity test (checking whether synthetic samples were distinguishable from the training set).

Lastly, we investigated the utility of the synthetic data in a downstream classification task. Specifically, a standard machine-learning classifier was trained exclusively on each synthetic dataset and tested on real held-out samples to measure classification accuracy, providing a real-world test of generative quality. All experiments were repeated on both the full and the reduced training sets to assess the effects of data scarcity on performance and generalizability.

### Technical implementation

All the models were implemented in Python (version 3.10.13) with PyTorch (version 2.5.1+cu124). The best configuration of each model was selected based on Balanced Energy Distance and was found using Weights & Biases sweeps, with a random search exploring both learning and architecture parameters. Training was conducted on FBK’s high-performance computing cluster Abacus with L40s and H100 Nvidia GPUs.

## Discussion

In earlier approaches, generative models are often evaluated with local fidelity metrics (such as the F1), offering limited insight into whether any model shortcomings are homogeneously distributed across the data manifold or confined to particular subpopulations, such as specific tissue types. However, in our tests we have found multiple architectures with high F1 scores dropping three-fold when considered in tissue-wise averages (e.g. from 0.9 to 0.3), as shown in Tables 1 and 2. In principle, these limitations could be mitigated by applying clustering procedures or systematic data partitioning. However, such methods add multiple layers of complexity and may inadvertently introduce bias, ultimately blurring the evaluation process.

**Table 1.** Performance metrics for the generative models on TCGA dataset. Average *±* standard deviation (5 generations).

|  | Full training set (8,124 samples) |  |  | Reduced training set (1,624 samples) |  |  |
| --- | --- | --- | --- | --- | --- | --- |
|  | GAN | DM | VAE | GAN | DM | VAE |
| balanced ED | 0.32 ± 0.0018 | 0.78 ± 0.020 | 2.57 ± 0.016 | 0.50 ± 0.028 | 0.72 ± 0.037 | 3.39 ± 0.050 |
| balanced PED | 0.93 ± 0.044 | 8.12 ± 0.30 | 24.11 ± 0.36 | 6.11 ± 0.95 | 5.19 ± 0.66 | 20.94 ± 0.28 |
| Adversarial ACC | 0.63 ± 0.0032 | 0.19 ± 0.0030 | 0.79 ± 0.0013 | 0.33 ± 0.0035 | 0.22 ± 0.011 | 0.90 ± 0.0033 |
| Classification ACC | 0.93 ± 0.0018 | 0.94 ± 0.0032 | 0.025 ± 0.0034 | 0.92 ± 0.00091 | 0.92 ± 0.0036 | 0.050 ± 0.013 |
| F1 score | 0.91 ± 0.0021 | 0.88 ± 0.0047 | 0.13 ± 0.0072 | 0.97 ± 0.0026 | 0.93 ± 0.0055 | 0.47 ± 0.014 |
| Tissue F1 | 0.93 ± 0.0020 | 0.89 ± 0.0050 | 0.13 ± 0.0045 | 0.98 ± 0.0025 | 0.95 ± 0.0054 | 0.16 ± 0.012 |
| Correlation | 0.99 ± 0.00044 | 0.98 ± 0.00035 | 0.87 ± 0.00075 | 0.97 ± 0.0055 | 0.96 ± 0.0020 | 0.80 ± 0.0030 |
| Frechet Score | 1.35 ± 0.11 | 4.30 ± 0.22 | 74.95 ± 0.65 | 11.40 ± 3.31 | 4.57 ± 0.24 | 101.04 ± 1.65 |

**Table 2.** Performance metrics for the generative models on GTEX dataset. Average *±* standard deviation (5 generations).

|  | Full training set (12,244 samples) |  |  | Reduced training set (2,448 samples) |  |  |
| --- | --- | --- | --- | --- | --- | --- |
|  | GAN | DM | VAE | GAN | DM | VAE |
| balanced ED | 0.24 ± 0.0059 | 0.93 ± 0.016 | 3.53 ± 0.023 | 0.32 ± 0.012 | 0.65 ± 0.030 | 5.61 ± 0.016 |
| balanced PED | 1.20 ± 0.15 | 7.91 ± 0.14 | 29.45 ± 0.21 | 0.63 ± 0.034 | 5.60 ± 0.18 | 63.90 ± 0.33 |
| Adversarial ACC | 0.64 ± 0.0031 | 0.24 ± 0.0027 | 0.91 ± 0.0019 | 0.31 ± 0.0065 | 0.28 ± 0.0092 | 0.90 ± 0.0014 |
| Classification ACC | 0.99 ± 0.0013 | 0.99 ± 0.0029 | 0.017 ± 0.0040 | 0.98 ± 0.0023 | 0.97 ± 0.00083 | 0.12 ± 0.0057 |
| F1 score | 0.93 ± 0.0019 | 0.94 ± 0.0023 | 0.28 ± 0.0082 | 0.99 ± 0.0013 | 0.97 ± 0.0017 | 0.18 ± 0.018 |
| Tissue F1 | 0.94 ± 0.0012 | 0.93 ± 0.0030 | 0.070 ± 0.0078 | 0.99 ± 0.00084 | 0.98 ± 0.0031 | 0.096 ± 0.018 |
| Correlation | 1.00 ± 0.00011 | 0.97 ± 0.0010 | 0.86 ± 0.00072 | 1.00 ± 0.00070 | 0.97 ± 0.0027 | 0.82 ± 0.0029 |
| Frechet Score | 2.50 ± 0.13 | 21.49 ± 0.48 | 176.55 ± 1.38 | 1.79 ± 0.13 | 8.28 ± 0.42 | 377.93 ± 4.10 |

By contrast, the adoption of statistical distances provides a clear and unified way to simultaneously assess the global and local alignment of generated and real data distributions. These distances are naturally extendable to include partial or localized neighborhood analyses and thus enable a balanced measure that weighs both global distribution alignment and neighborhood fidelity. In practice, we found that simply optimizing the balanced energy distance was sufficient to attain strong local performance, reflected by high tissue-level F1 scores, without requiring explicit modeling of these subgroups.

No substantial advantage was observed in using the pointwise empirical distance over the energy distance for these particular *omics* datasets. Both measures selected similar architectures and yielded comparable rankings throughout training (Tables 1 and 2). One likely explanation for this equivalence is that the dataset in question could be easily normalized and presented moderate noise levels (at most), reducing the value of manifold evaluation and noise reduction that the PED offers.

A further benefit of relying on statistical distances was observed during the training phase of generative adversarial networks, where the default loss functions are often not directly interpretable. Tracking the statistical distances provides an objective, smoothly varying indicator of the gap between synthetic and real data distributions, as highlighted in the rightmost panels of Figure 1 for ED (at the top) and PED (at the bottom). This perspective becomes all the more valuable when combined with a thorough comparison on the validation set, specifically through a twofold test for likelihood (whether synthetic data could plausibly come from the real distribution; leftmost panels, Figure 1 and identity (ensuring that the synthetic data are distinguishable from the original training set; middle panels, Figure 1. Such tests guard against overfitting, where a model might memorize training samples rather than learn a more general underlying distribution. Our experiments confirm that only certain architectures succeed in passing both the likelihood and identity tests. The same architectures also generate synthetic data that supports downstream classification tasks at levels matching those achieved when training directly on real data. This alignment between robust distributional fidelity (as demonstrated by statistical distances) and consistent downstream utility (as evidenced by classification performance) underscores the value of these distance-based criteria as reliable indicators of model fitness.

**Fig. 1:**
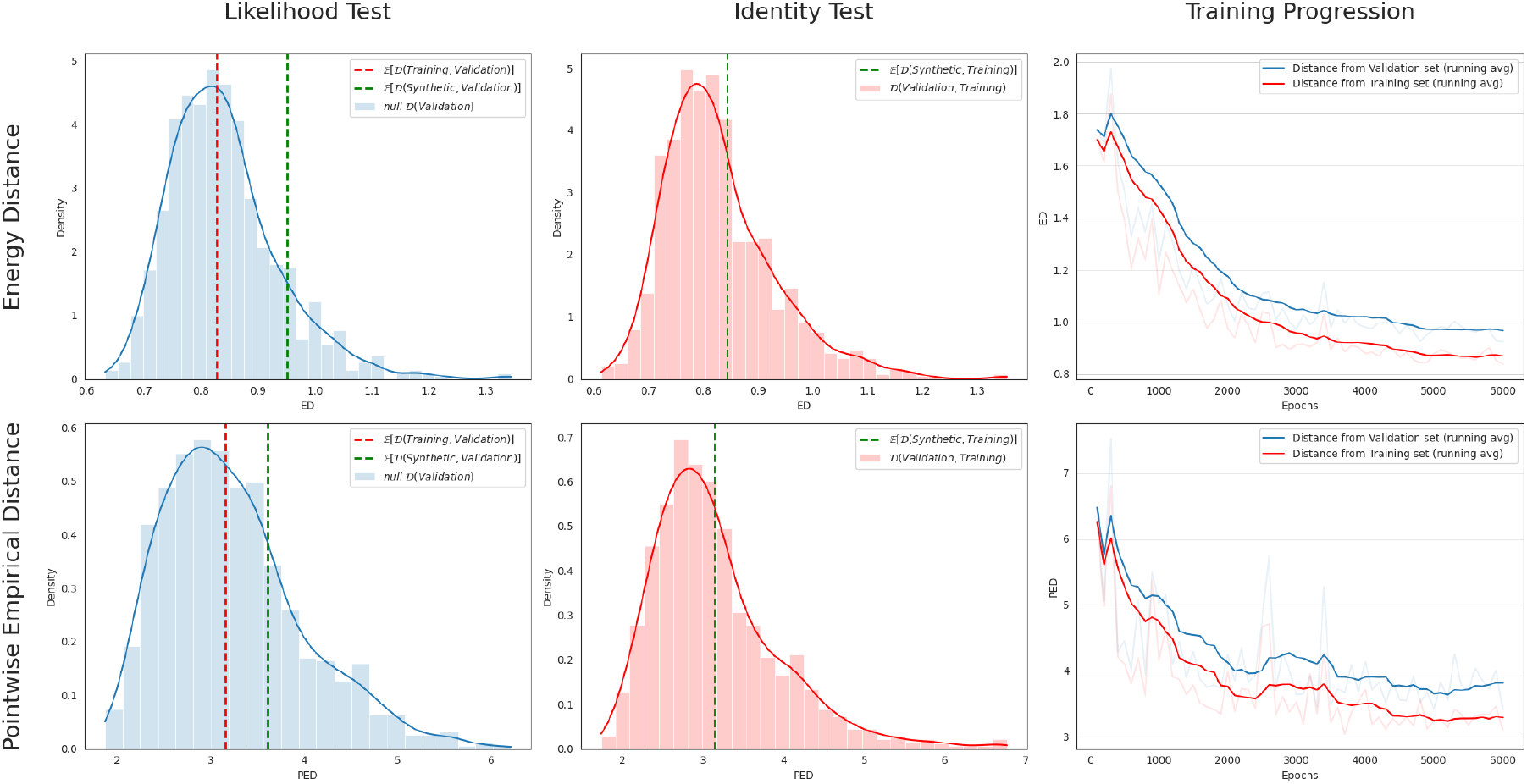
Our proposed framework for the evaluation of generative models: Likelihood Test, Identity Test, and Training Progression measured with Energy Distance and Pointwise Empirical Distance. The plots show the results for a WGAN-GP trained on GTEx data.

We believe these findings to be particularly relevant in domains such as healthcare and molecular biology, where datasets exhibit high manifold complexities and comparatively low sample size. The ability to measure both global and local alignment of synthetic to real data in an unbiased, model-agnostic way makes these statistical distances a powerful tool, also helping overcome the inherent limitations of classical local fidelity metrics such as F1 (Powers, 2020). The adoption of statistical distances can help guide the optimization of generative models to balance realism, diversity, and privacy requirements while maintaining interpretability. We believe these distances will become increasingly central in guiding future progress in generative modeling, from data augmentation in resource-limited settings to more advanced applications where noise and manifold complexity pose considerable challenges.

## Author contributions statement

NL conceived the experiments on statistical distances, performed their implementation, interpreted the results, and substantially contributed to writing the paper. RM built the overall pipeline for generative model validation, comparing statistical distances with alternative metrics, and developed the Generative Adversarial Network code. RM also prepared the datasets and contributed to writing the relevant sections and interpreting the results. GL and FR assisted in data preparation and implemented the Stable Diffusion and Variational Autoencoder architectures, writing their respective sections. SB, MC contributed to designing the overall study structure, drafted the first version of the manuscript, and wrote parts of the introduction. MC, GJ supervised the project and provided conceptual advice. GJ secured funding and provided direction for the entire study. All authors contributed to writing and revising the final manuscript.

## Acknowledgments and Funding

The results shown here are in whole or part based upon data generated by the TCGA Research Network^1^. This work is partially funded by the EU through the 3DSecret project under the HORIZON-EIC-2022-PATHFINDER-OPEN-01-01 programme (grant no. 101099066) and NextGenerationEU - National Recovery and Resilience Plan (PNRR) — Research program CN00000013 “National Centre for HPC, Big Data and Quantum Computing”, Spoke 8: Insilico Medicine and Omics Data (CN 00000013 — Avviso n. 3138 del 16 dicembre 2021).

## Footnotes

1 https://www.cancer.gov/tcga

